# DeepMF: Deciphering the Latent Patterns in Omics Profiles with a Deep Learning Method

**DOI:** 10.1101/744706

**Authors:** Lingxi Chen, Jiao Xu, Shuai Cheng Li

**Affiliations:** City University of Hong Kong, 83 Tat Chee Ave, Kowloon Tong, HongKong, China

## Abstract

With recent advances in high-throughput technologies, *matrix factorization* techniques are increasingly being utilized for mapping quantitative omics profiling matrix data into low-dimensional embedding space, in the hope of uncovering insights in the underlying biological processes. Nevertheless, current matrix factorization tools fall short in handling noisy data and missing entries, both deficiencies that are often found in real-life data. Here, we propose DeepMF, a deep neural network-based factorization model. DeepMF disentangles the association between molecular feature-associated and sample-associated latent matrices, and is tolerant to noisy and missing values. It exhibited feasible subtype discovery efficacy on mRNA, miRNA, and protein profiles of medulloblastoma cancer, leukemia cancer, breast cancer, and small-blue-round-cell cancer, achieving the highest clustering accuracy of 76%, 100%, 92%, and 100% respectively. When analyzing data sets with 70% missing entries, DeepMF gave the best recovery capacity with silhouette values of 0.47, 0.6, 0.28, and 0.44, outperforming other state-of-the-art MF tools on the cancer data sets Medulloblastoma, Leukemia, TCGA BRCA, and SRBCT. Its embedding strength as measured by clustering accuracy is 88%, 100%, 84%, and 96% on these data sets, which improves on the current best methods 76%, 100%, 78%, and 87%. DeepMF demonstrated robust denoising, imputation, and embedding ability. It offers insights to uncover the underlying biological processes such as cancer subtype discovery. Our implementation of DeepMF can be found at https://gitlab.deepomics.org/jiaox96/DeepMF.

## Introduction

Recent advances in high-throughput technologies have eased the quantitative profiling of biological data and enabled many *in silico* studies to elucidate complex biological processes [1]. In many cases, the biological data are captured in a matrix with molecular features such as gene, mutation locus, or species as rows and samples/repetition as columns. Values in the matrices are typically measurements such as expression abundances, mutation levels, or species counts. Based on the assumption that samples with similar phenotype (or molecular features) that participate in a similar biological process will share similar distribution of biological variation [1], patterns shared by a significant number of entries in these matrices may yield insights on important biological processes.

Clustering through methods like *k*-means and hierarchical clustering have been used to identify these patterns [2, 3]. The success of these studies are contingent on the ability of these clustering methods to capture the underlying structures or models of the interaction patterns.

Matrix factorization (MF), as given by the formula ***A***_*M*×*N*_ ≈ ***U***_*M*×*K*_ × ***V*** _*K*×*N*_ in **Figure 1A**, is a popular approach to the problem. Numerous researches have applied MF to identify latent structures (***U*** and ***V***) from a given matrix of biological data (***A***_*M*×*N*_). A good factorization technique would ensure that as much information as possible from ***A*** is conserved [1, 2]. Here, we hope to uncover *K* hidden signatures from the complex biological processes. We refer to ***U*** as the molecular feature latent matrix, since the values in each column of ***U*** are continuous weights illustrating the relative participation of a molecule in each inferred biology process signature. We call ***V*** the sample latent matrix, as each row of ***V*** depicts the fractions of samples in the matched biological process signature. Molecular features or sample subgroups can be detected by finding patterns in the molecular feature latent matrix and sample latent matrix, respectively. MF has been successfully applied to multiple data modalities [1]. For instance, it has been used to detect leukemia cancer subtype based on expression profiles. By combining gene expression and DNA methylation data, MF has been used to classify HPV subtypes in head and neck tumors [4]. It has also been used to define COSMIC mutational signatures in pan-cancer studies [5–7].

**Figure 1.**
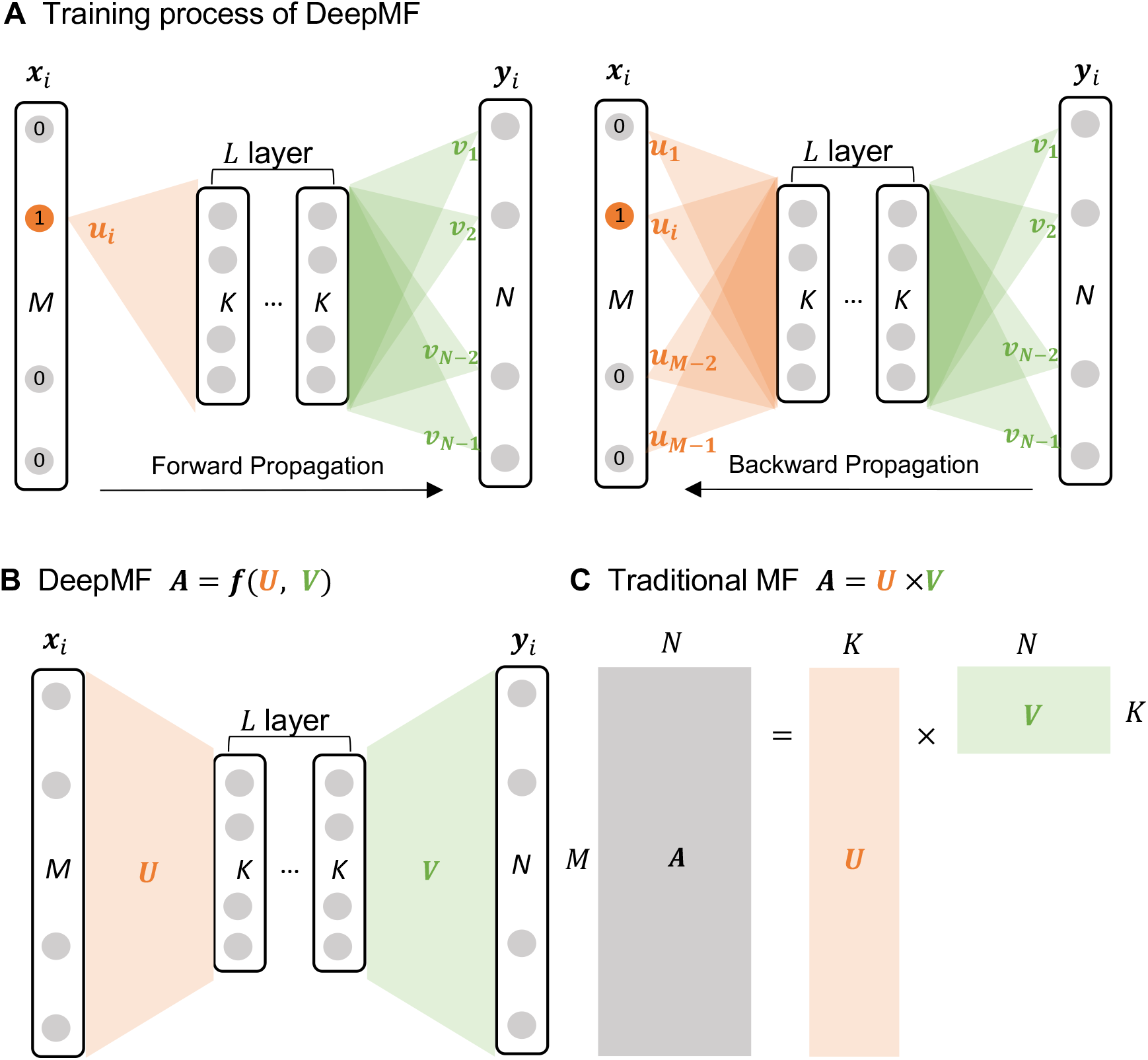
DeepMF Structure Overview (A) The training process of DeepMF. The rectangle represents the *K* dimensional gene latent vector ***u*** ∈ ℝ^*K*^ or sample latent vector ***v*** ∈ ℝ^*K*^. (B-C) Illustration of DeepMF and MF, respectively.

MF methods, such as Principal Component Analysis (PCA), Independent Component Analysis (ICA), and Non-Negative Matrix Factorization (NMF), are widely used to extract the low-dimensional latent structure from high-dimensional biological matrix [1]. Intuitively, PCA finds governing variation in high-dimensional data, securing the most important biological process signatures that differentiate between samples [8]. ICA separates mixed signal matrix into statistically independent biological process signatures [9]. NMF-based approaches extracted latent matrices with non-negative constraints [10, 11].

Despite the effectiveness of MF in interpreting biological matrices, several limitations persist in practice. First, real-world data are often plagued with many types of noises, e.g. systematic noise, batch effect, and random noise [12], which potentially mask signals in the downstream process. Second, high throughput omics data frequently suffer from missing values due to various experimental settings [13], whereas the majority of MF tools have no support for input matrix with missing values. At present, the standard practice to deal with these two problems is to perform denoising and imputation prior to MF. However, even when these problems are mitigated, the MF techniques mentioned would still be unable to uncover any non-linear relationship, since they assume a linear association between molecular feature latent variables and sample latent variables.

In this work, we propose a deep neural network-based matrix factorization framework, DeepMF (**Figure 1 B**), which learns the non-linear association between molecular feature latent matrix and sample latent matrix, tolerant with noisy and missing entries. DeepMF demonstrated robust denoising, imputation, and embedding ability in simulated instances. It outperformed the existing MF tools on subtype discovery in omics profiles of medulloblastoma cancer, leukemia cancer, breast cancer, and small-blue-round-cell cancer, with the highest clustering accuracy on all the four datasets collected for this work. Furthermore, with 70% data randomly removed, DeepMF demonstrated the best recovery capacity with silhouette values 0.47, 0.6, 0.28, and 0.44. It also displayed the best embedding power on the four datasets, with clustering accuracy of respectively 88%, 100%, 84%, and 96%, which improves on the current best methods 76%, 100%, 78%, and 87%.

## Background

Recent advances in high-throughput technologies have eased the quantitative profiling of biological data and enabled many *in silico* studies to elucidate complex biological processes [1]. In many cases, the biological data are captured in a matrix with molecular features such as gene, mutation locus, or species as rows and samples/repetition as columns. Values in the matrices are typically measurements such as expression abundances, mutation levels, or species counts. Based on the assumption that samples with similar phenotype (or molecular features) that participate in a similar biological process will share similar distribution of biological variation [1], patterns shared by a significant number of entries in these matrices may yield insights on important biological processes. Clustering through methods like *k*-means and hierarchical clustering have been used to identify these patterns [2, 3]. The success of these studies are contingent on the ability of these clustering methods to capture the underlying structures or models of the interaction patterns.

Matrix factorization (MF), as given by the formula ***A***_*M*×*N*_ ≈ ***U*** _*M*×*K*_ × ***V*** _*K*×*N*_ in **Figure 1A**, is a popular approach to the problem. Numerous researches have applied MF to identify latent structures (***U*** and ***V***) from a given matrix of biological data (***A***_*M*×*N*_). A good factorization technique would ensure that as much information as possible from ***A*** is conserved [1, 2]. Here, we hope to uncover *K* hidden signatures from the complex biological processes. We refer to ***U*** as the molecular feature latent matrix, since the values in each column of ***U*** are continuous weights illustrating the relative participation of a molecule in each inferred biology process signature. We call ***V*** the sample latent matrix, as each row of ***V*** depicts the fractions of samples in the matched biological process signature. Molecular features or sample subgroups can be detected by finding patterns in the molecular feature latent matrix and sample latent matrix, respectively. MF has been successfully applied to multiple data modalities [1]. For instance, it has been used to detect leukemia cancer subtype based on expression profiles. By combining gene expression and DNA methylation data, MF has been used to classify HPV subtypes in head and neck tumors [4]. It has also been used to define COSMIC mutational signatures in pan-cancer studies [5–7].

MF methods, such as Principal Component Analysis (PCA), Independent Component Analysis (ICA), and Non-Negative Matrix Factorization (NMF), are widely used to extract the low-dimensional latent structure from high-dimensional biological matrix [1]. Intuitively, PCA finds governing variation in high-dimensional data, securing the most important biological process signatures that differentiate between samples [8]. ICA separates mixed signal matrix into statistically independent biological process signatures [9]. NMF-based approaches extracted latent matrices with non-negative constraints [10, 11].

Despite the effectiveness of MF in interpreting biological matrices, several limitations persist in practice. First, real-world data are often plagued with many types of noises, e.g. systematic noise, batch effect, and random noise [12], which potentially mask signals in the downstream process. Second, high throughput omics data frequently suffer from missing values due to various experimental settings [13], whereas the majority of MF tools have no support for input matrix with missing values. At present, the standard practice to deal with these two problems is to perform denoising and imputation prior to MF. However, even when these problems are mitigated, the MF techniques mentioned would still be unable to uncover any non-linear relationship, since they assume a linear association between molecular feature latent variables and sample latent variables.

In this work, we propose a deep neural network-based matrix factorization framework, DeepMF (**Figure 1 B**), which learns the non-linear association between molecular feature latent matrix and sample latent matrix, tolerant with noisy and missing entries. DeepMF demonstrated robust denoising, imputation, and embedding ability in simulated instances. It outperformed the existing MF tools on subtype discovery in omics profiles of medulloblastoma cancer, leukemia cancer, breast cancer, and small-blue-round-cell cancer, with the highest clustering accuracy on all the four datasets collected for this work. Furthermore, with 70% data randomly removed, DeepMF demonstrated the best recovery capacity with silhouette values 0.47, 0.6, 0.28, and 0.44. It also displayed the best embedding power on the four datasets, with clustering accuracy of respectively 88%, 100%, 84%, and 96%, which improves on the current best methods 76%, 100%, 78%, and 87%.

## Method

### Matrix Factorization by Deep Neural Network

In this section, we introduce the DeepMF architecture and the loss function used for its training. Unless stated otherwise, symbols in bold font refer to vectors or matrices.

### Matrix Factorization

In **Figure 1C**, assume the input matrix ***A*** is of dimension *M* × *N*, where *M* is the number of features, and *N* is the number of samples. A row represents a feature while a column represents a sample or a replication. The element ***A***_*ij*_ refers to the measured values for feature *F*^*i*^ on sample *S*^*j*^, 1 ≤ *i* ≤ *M*, 1 ≤ *j* ≤ *N*.

Matrix factorization assumes the dot product of feature latent factor ***u***^*i*^ and sample latent factor ***v***^*j*^ to capture the interactions between feature *F*^*i*^ and sample *S*^*j*^, where ***u***^*i*^ and ***v***^*j*^ are vectors of size *K* which encode structures that underlie the data; that is, the predicted element of feature *F*^*i*^ on sample *S*^*j*^ is calculated as:

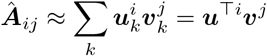

The predicted matrix 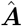 can be thought of as the product of the feature latent factor matrix ***U*** and sample latent factor matrix ***V***, 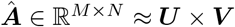, where ***U*** ∈ ℝ^*M*×*K*^, **V** ∈ ℝ^*K*×*N*^, *K* ≪ *M, N*. The objective function is 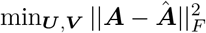.

### Framework architecture

DeepMF is modeled to learn the complex interactions between two latent factors ***U*** and ***V*** based on neural network. **Figure 1B** illustrates the network architecture of DeepMF. The input layer has *M* neurons, corresponding to *M* features in the matrix. The output layer has *N* nodes to model the *N* column samples. DeepMF is to capture the non-linear interaction between ***U*** and ***V***. As illustrated in **Figure 1B**, the network utilizes *L* hidden layers of *K* nodes each. All the nodes in the hidden layers are fully connected and paired with `ReLU` activation function. The number of nodes, *K*, corresponds to the dimensionality of the latent space in matrix factorization. The network is be sufficiently complex in order to approximate *f*(***U***, ***V***), a non-linear function for the interaction between feature and sample latent factors.

### Training

The matrix ***A*** ∈ ℝ^*M*×*N*^ contains *M* features. Each feature *F*^*i*^ corresponds to one input data point ***x***^*i*^ ∈ ℝ^*M*^ and output label ***y***^*i*^ ∈ ℝ^*N*^, where ***x***^*i*^ is one-hot encoded and ***y***^*i*^ is the *i*-th row of matrix ***A***.

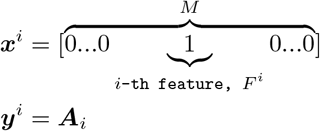

The loss function consists of two parts, one for global trends and one for local trends. For a pair of feature *F*^*i*^ and sample *S*^*j*^, *global proximity* refers to the proximity between real measurement ***A***_*ij*_ and predicted value 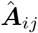. The preservation of global proximity is fundamental in matrix factorization. On the other hand, if two samples possess many common features, they tend to be similar. We refer to this similarity as *sample local proximity*. We define *feature local proximity* similarly. By introducing these local proximities into the loss function, we aim to identify and preserve the sample-pairwise and feature-pairwise structures in the low-dimensional latent space.

For global proximity, we minimize the 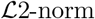 of the residual:

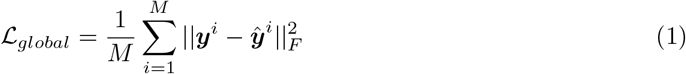

For the local proximities, we use feature local proximity 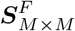 and sample local proximity 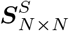 as supervised information. They respectively constrain the similarity of the latent representations of features and samples. Given matrix ***A***_*M*×*N*_, we obtain the feature similarity matrix 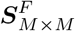 and sample similarity matrix 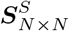 as

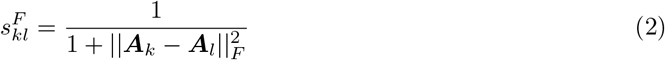

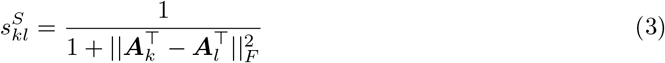

where ***A***_*k*_ and ***A***_*l*_ refer to the *k*-th and *l*-th row of matrix 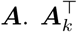 and 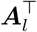 refer to the *k*-th and *l*-th column of matrix ***A***.

With ***S***^*F*^ and ***S***^*S*^, we define 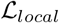 to preserve the local proximity of learned latent matrices ***U*** and *V* :

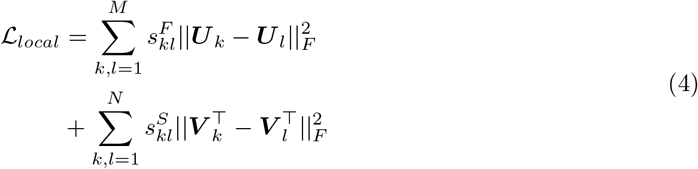

where ***U***_*k*_ and ***U***_*l*_ refer to the *k*-th and *l*-th row of feature latent matrix ***U***, 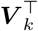 and 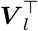 refer to the *k*-th and *l*-th column of sample latent matrix ***V***.

The objective function 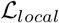 incurs a penalty when similar features and similar samples are embedded far away in the latent space. Hence, two features or samples with low similarity will be driven nearer in the embedding space. To prevent this, we first identify the remote sample-sample or feature-feature pair from feature and sample local proximity matrices by *k*-means. Then we mark their local similarity to zero to exclude them from 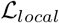 constraints.

To avoid overfitting and constrain the latent matrices ***U*** and ***V***, an 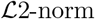 regularization is incorporated with ***U***, ***V***, and model hidden layer weights ***W***_*hidden*_.

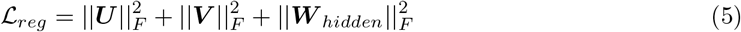

Our final loss function incorporates all the above constraints, with two additional hyperparameters *α* and *β*, as follows:

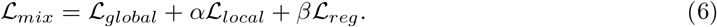

### Dealing with missing value

To be tolerant to missing values, DeepMF discards the missing entries in back-propagation by a variational 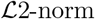. Denote *ξ* as a missing value.

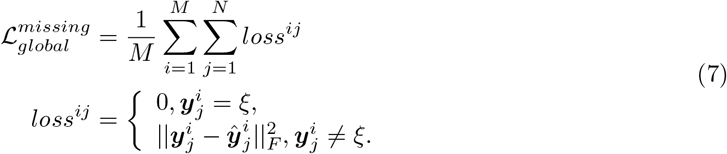

Then, DeepMF can infer a missing value ***A***_*αβ*_ by utilizing the trained model

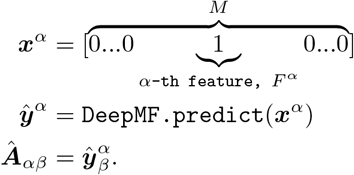

### DeepMF architecture parameter selection

If the data assumes *C* (*C* ≥ 2) clusters with respect to samples, we recommend that the network structure be pruned as guided by the validation loss 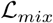 in the range of *K* ∈ [2, *C*] and *L* ∈ [1, +*∞*). For a matrix ***V***_*K*×*N*,(*K*<*N*)_, a rank of *C* is enough to represent the latent hierarchical structure for a *C*-clustering problem, thus *K* ≤ *C*. To extract simple patterns like the linear association between feature and sample, *L* = 1 suffices. A larger *L* would provide more complexity in the latent space of DeepML. For hyperparameter tuning, we recommend running each *K, L* combination more than ten times with different random weights initialization to avoid possible local optima.

### DeepMF interpretation

DeepMF operates on the basis of the ML formula 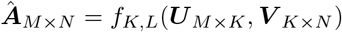, where *f*_*K,L*_ refers to a collection of non-linear mapping, ***U*** is the feature latent factor matrix, and ***V*** is the sample latent factor matrix (**Figure 1**). It learns about missing values in training, and imputes them in prediction. Since DeepMF is trained by minimizing the loss between ***A*** and 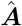, denoising is built into the learning process.

The two extracted matrices ***U*** and ***V*** are modeled to uncover the underlying latent structures of the features and samples, respectively. They can hence be applied to features and samples related clustering and pattern recognition tasks for data interpretation.

### Simulation data generation

To evaluate DeepMF, we simulated three patterns, each which consists of matrices of sizes 1000 × 600, 10 × 6, and 100 × 60 as shown in **Figures 2, S1**, and **S2**. Then, we randomly removed 10%, 50%, and 70% of the matrices to make them sparse.

**Figure 2.**
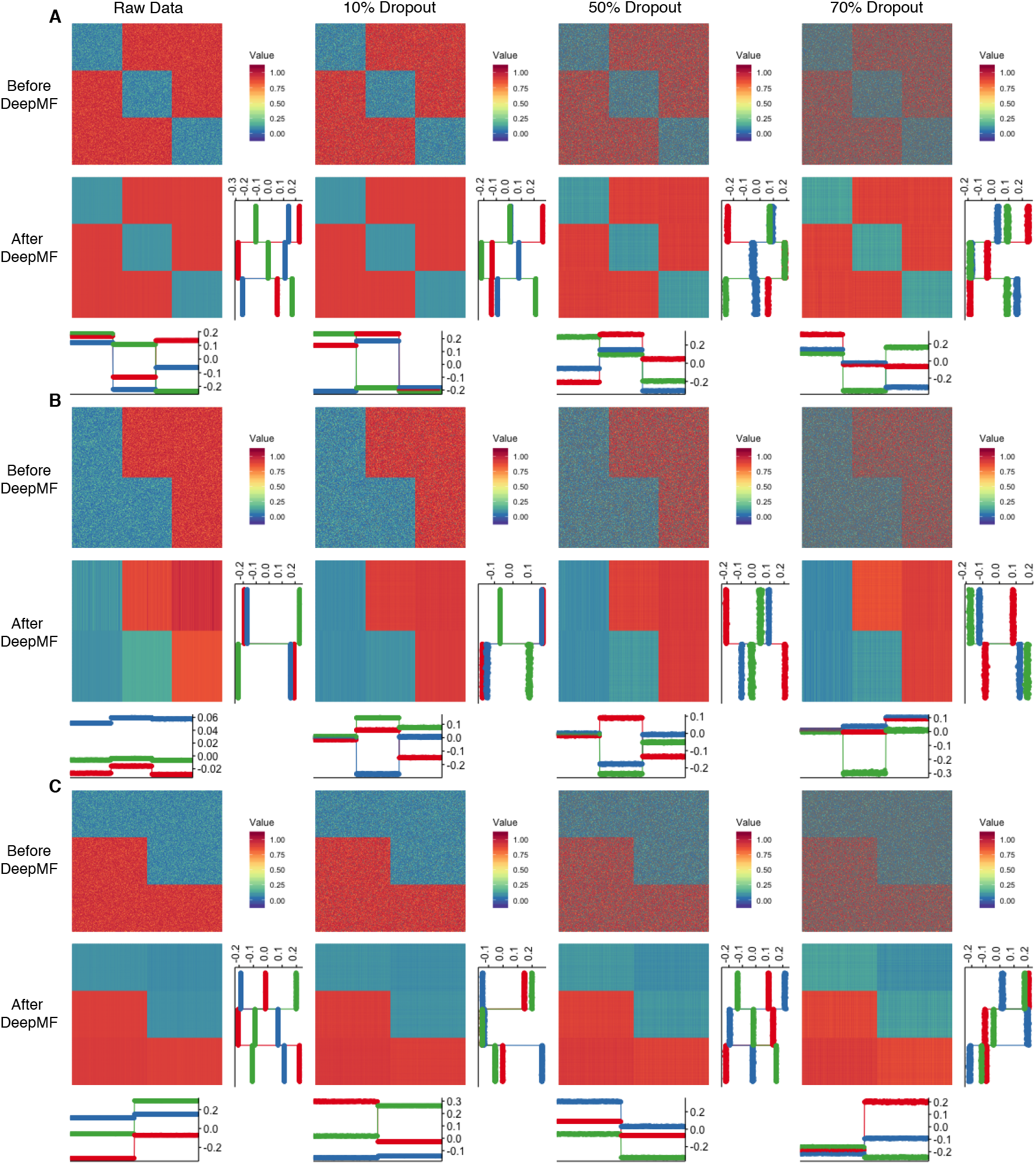
DeepMF performance on 1000 × 600 synthetic matrices DeepMF denoising, imputation, and factorization performance on 1000 × 600 synthetic matrices with different pattern. Inside each pattern, from left to right: raw matrix, 10% random dropout, 50% random dropout, 70% random dropout; from top to bottom: before DeepMF, and DeepMF. The horizontal line plot show the sample latent factors, the vertical line plot refer to feature latent factors. **A** Matrix with pattern A **B** Matrix with pattern B **C** The transpose matrix of pattern B

### Cancer subtyping experiments

For real datasets, the four cancer datasets as follows are used.

### Cancer data preparation

#### Medulloblastoma data set

Gene expression profiles from childhood brain tumors medulloblastomas were obtained from Brunet’s work [2]. It consists of classic and desmoplastic subtypes of size 25 and 9, respectively. We further extracted the top 100 differentially expressed genes using “limma” R package [14].

#### Leukemia data set

The Leukemia data set was obtained from R package “NMF” with the command “`data(esGolub)`” [10]. It stores Affymetrix Hgu6800 microarray expression data from 38 Leukemia cancer patients, where 19 patients with B cell Acute Lymphoblastic Leukemia (B-cell ALL), eight patients with T cell Acute Lymphoblastic Leukemia (T-cell ALL), as well as 11 patients with Acute Myelogenous Leukemia (AML). The 236 most highly diverging genes were selected by comparison on their coefficient of variation using limma R package [14].

#### TCGA BRCA data set

A subset of human breast cancer data generated by The Cancer Genome Atlas Network (TCGA) was obtained from R package mixOmics [15]. It holds 150 samples with three subtypes Basal-like, Her2, and LumA, of size 45, 30 and 75, respectively. The top 55 correlated mRNA, miRNA, and proteins which discriminate the breast cancer subtypes subgroups Basal, Her2, and LumA were selected using the mixOmics DIABLO model.

#### SRBCT data set

The Small Round Blue Cell Tumors (SRBCT) data set holds the expression profiles of the top 96 ranked genes [16]. It contains 63 samples of four classes, Burkitt Lymphoma (BL, 8 samples), Ewing Sarcoma (EWS, 23 samples), Neuroblastoma (NB, 12 samples), and Rhabdomyosarcoma (RMS, 20 samples). The processed and normalized data were acquired from the R mixOmics package [15].

### Decomposition baselines

We compared the decomposition efficacy on DeepMF against four methods, PCA (FactoMineR [17]), ICA (fastICA [18]), Bayesian-based NMF (CoGAPS [11]), and gradient-based NMF (NMF [10]). We fit all model with log-treated matrices. All tools were executed with their recommended settings; that is, `prcomp` function in package “FactoMineR”; fastICA with algorithm type “parallel”, function “logcosh”, alpha 1, method “R”, row normalization 1, maxit 200, tol 0.0001; CoGAPS with 5000 iterations; NMF with method “brunet” and 200 runs.

As CoGAPS and NMF accept only non-negative values, we used `NMF.posneg` to transform the input matrices into corresponding non-negative matrices.

### Imputation baselines

We evaluated the DeepMF imputation efficiency by comparing it with two popular imputation approaches, MeanImpute, and SVDImpute.

#### MeanImpute

MeanImpute adopted the approach that the missing entries are to be substituted by the mean of the current values of a particular feature in all samples. We used the mean impute function in the R package “CancerSubtypes”.

#### SVDImpute

SVDImpute first centers the matrix, replaces all missing values by 0, decomposes the matrix into the eigenvectors. Then, SVDImpute predicts the NA values as a linear combination of the *k* most significant eigenvectors [19]. We chose SVDImpute as an imputation baseline since the mechanism behind it is similar to DeepMF. The *k* most significant eigenvectors can be analogized to the *k*-dimensional latent matrix in DeepMF. We used R package “pcaMethods” in practice.

### Evaluation Metrics

#### Silhouette width

The silhouette width measures the similarity of a sample to its class compared to other classes [20]. It ranges from −1 to 1. A higher silhouette value implies a more appropriate clustering. A silhouette value near 0 intimates overlapping clusters, and a negative value indicates that the clustering has been performed incorrectly.

We adopted the silhouette width to evaluate the model’s denoising and imputation power. We used the ground-truth subtype classes as the input cluster labels. Then, the silhouette width for a given matrix was calculated with Euclidean distance using the R package “cluster”.

#### Adjusted Rand Index

We also used the adjusted Rand index to evaluate the clustering accuracy. The adjusted Rand index measures the similarity between predicted clustering results and actual clustering labels [21]. A value close to 0 indicates random labeling, and a value of 1 demonstrates 100% accuracy of clustering.

To check the cancer subtyping effectiveness of different matrix factorization tools. We first used the R hierarchy clustering packaging “hclust” to obtain the sample latent factor matrices in order to partition samples into subgroups, through the Euclidean distance and “ward.D2” linkage. Then, we computed the adjusted Rand index to measure the clustering accuracy via the R package “fpc”

## Results

### Denoising, imputation, and embedding evaluation on synthetic data

To evaluate the denoising, imputation, and embedding efficacy of DeepMF, we first generated three patterns A, B and C, each which consists of matrices of size 1000 × 600, 10 × 6, and 100 × 60 in (**Figures 2, S1, S2**). Matrices with pattern A hold three subgroups in feature and sample. Pattern B has two subgroups in feature and three subgroups in sample. Pattern C matrices are transposed of pattern B of dimension 600 × 1000, 6 × 10, and 60 × 100. Then we generated sparse matrices randomly by dropping the entries of matrices with rate 10%, 50%, and 70%.

**Figures 2, S1, S2** show the performance of DeepMF on the raw matrix and sparse matrix with size 1000 × 600, 10 × 6, and 100 × 60, respectively. In **Figures 2 A, S1 A, S2 A**, the DeepMF predicted matrices significantly reduced the noisy and missing entries. In spite of the noise and 70% missing entries, the feature latent factors and sample latent factors generated by DeepMF consistently uncovered ground truth feature subgroups and sample subgroups with 100% accuracy. The same conclusion applies to pattern B and pattern C (**Figures 2 B-C, S1 B-C, S2 B-C**). We note that pattern B matrices and pattern C matrices are transposed, which suggests that DeepMF can uncover the feature and sample subclasses either from a feature-sample matrix or its transposed matrix. Since fitting a matrix with *N* < *M* is more efficient than a matrix with *N* > *M* in DeepMF, it may be beneficial to do so, as long as it is unnecessary to adhere to the paradigm of “treating the feature as row and sample as column” [1].

### DeepMF accurately elucidates cancer subtypes on multiple cancer omics data sets

To discover complex biological processes from massive amounts of high-throughput matrix data, researchers customarily separate features or samples with similar profiles into biologically significant partitions, with the assistance of clustering or pattern recognition techniques. Here, to demonstrate how DeepMF can assist in this biological discovery, we collected a series of cancer omics data sets, namely the Medulloblastoma data set (mRNA) [2], Leukemia data set (mRNA) [2, 10], TCGA BRCA data set (mRNA, miRNA, protein) [15], and small blue round cell tumor (SRBCT) data set (mRNA) [15, 16]. Then, we employed them as benchmark sets for cancer subtyping analysis.

We first verified the correctness of the output matrices. **Figure 3** shows that DeepMF reduced the noise in raw matrices while preserving cancer subtype structures on all cancer omics data sets. Silhouette validation corroborated that the in-cluster similarity and out-cluster separation were enhanced after DeepMF processing; that is, the average silhouette value was increased from 0.26 to 0.56 for Medulloblastoma data set, from 0.35 to 0.66 for Leukemia data set, from 0.19 to 0.47 for TCGA BRCA data set, from 0.31 to 0.58 for SRBCT data set, respectively (see **Figure 3**).

**Figure 3.**
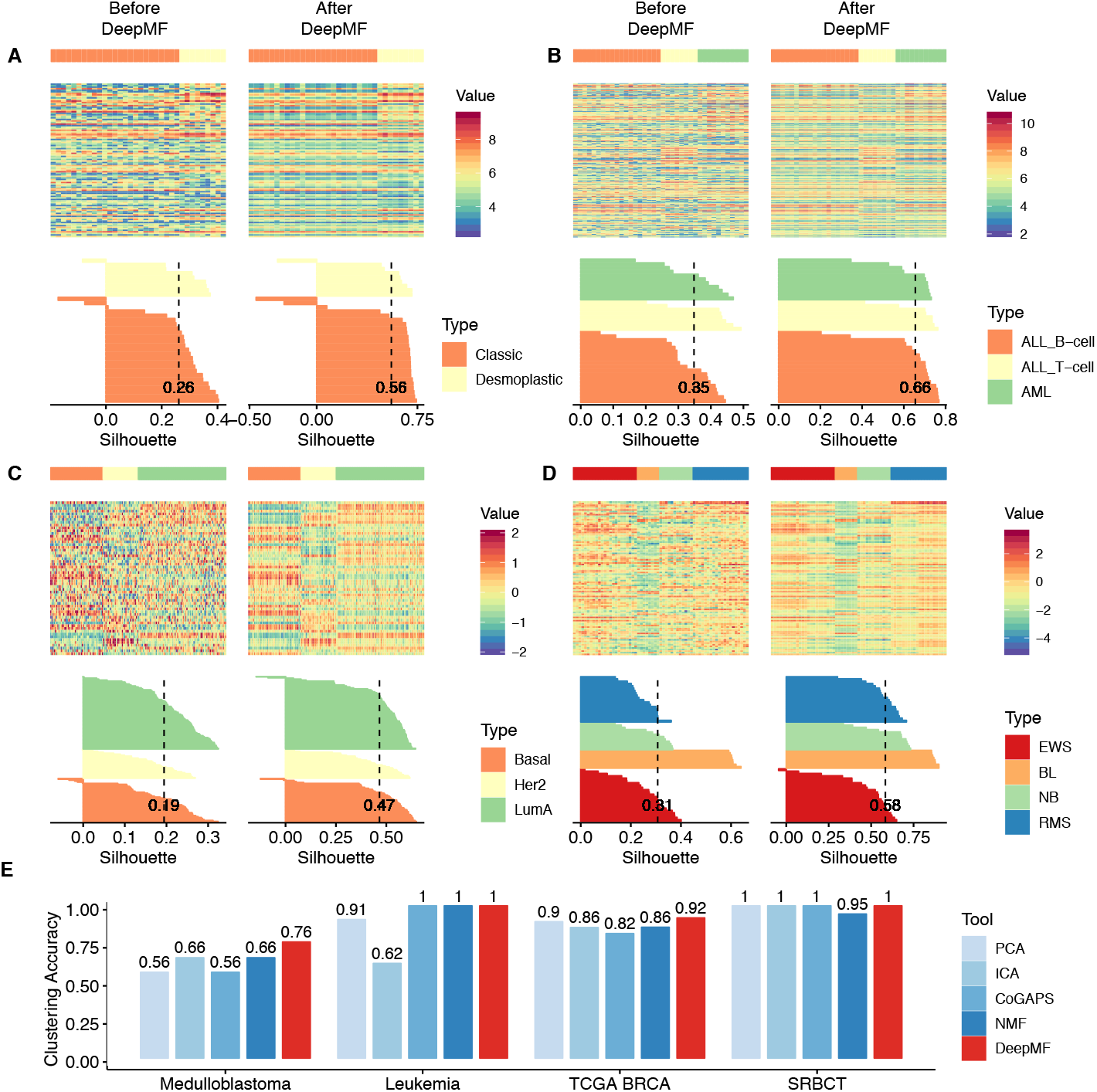
DeepMF denoising and factorization on cancer data sets **A-D** The heatmap presentation and Silhouette width of four cancer data sets. Left: before DeepMF. Right: after DeepMF. **A** Medulloblastoma data set **B** Leukemia data set **C** TCGA BRCA data set **D** SRBCT data set **E** Clustering accuracy of cancer subtyping on sample latent matrices generated by five matrix factorization tools on different cancer data sets.

We then checked whether the DeepMF produced sample latent matrix preserves the cancer subtype information. We compared the decomposition efficiency on DeepMF against four traditional matrix factorization methods, PCA (FactoMineR [17]), ICA (fastICA [18]), Bayesian-based NMF (CoGAPS [11]), and gradient-based NMF (NMF [10]). We fitted high dimensional raw matrices into DeepMF and the above four tools, extracted the low-dimensional sample latent matrices with rank *K* = 2 for Medulloblastoma data set, rank *K* = 3 for Leukemia data set, rank *K* = 3 for TCGA BRCA data set, and rank *K* = 4 for SRBCT data set, respectively. The DeepMF structure configuration in training is listed in **Table S1**. To escape from local optima caused by DeepMF random weight initialization, we conducted ten different runs for each data set configuration and selected the latent matrices with minimal loss. Next, we applied hierarchical clustering into obtained sample latent matrices (**Figure S3**). Clustering accuracy is evaluated by the adjusted Rand index, which measures the overlap between the inferred clusters and ground-truth subtype, a score of 0 signifies random labeling and 1 denotes perfect inference. In **Figure 3**, DeepMF outperforms all four methods and manifests the best embedding strength, with highest clustering accuracy of 76% for Medulloblastoma data set, 92% for TCGA BRCA data set, and 100% accuracy for Leukemia and SRBCT data sets.

### DeepMF captures the cancer subtype patterns despite 70% random dropouts

Several studies have suggested that missing values in large-scale omics data can drastically obstruct the interpretation of complex biological processes, such as unsupervised cancer subtyping [22]. At present, this is most commonly treated by imputing the missing values before performing downstream analysis of multi-omics data. To evaluate the imputation efficiency of DeepMF, we randomly dropout 70% entries on all four cancer data sets, then fit the sparse matrices into DeepMF and two imputation baselines: MeanImpute and SVDImpute. We selected MeanImpute by considering its popularity. From the perspective of imputation mechanism, we can regard SVDImpute as a linear analogy of DeepMF. The DeepMF structure configuration in training is listed in **Table S1**. To avoid local optima, we conducted ten different runs for each data set configuration and picked the one with minimal loss. **Figure 4** demonstrates that for all 70% missing rate data sets, both DeepMF and SVDImpute recovered distinctive cancer subtype structures, while the MeanImpute approach was unable to reconstruct a clearly visible pattern. Silhouette validation confirmed that DeepMF reduced the most substantial interior cluster heterogeneity and out-cluster similarity, with the largest average silhouette value of 0.47 for the Medulloblastoma data set, 0.6 for the Leukemia data set, 0.28 for TCGA BRCA data set, and 0.44 for SRBCT data set.

**Figure 4.**
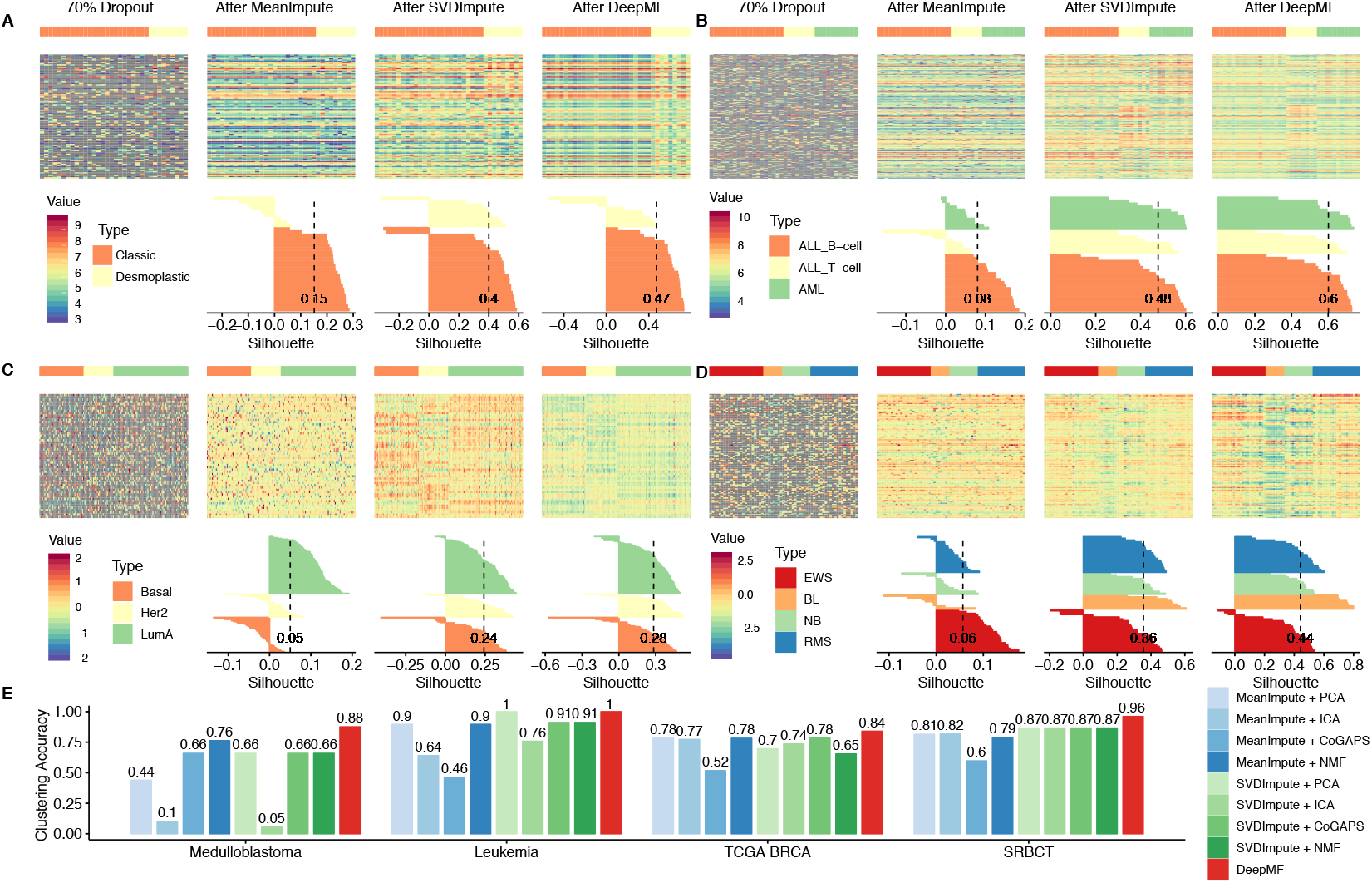
DeepMF’s imputation and factorization effect on cancer data sets with 70% random dropout **A-D**The heatmap presentation and Silhouette width of four cancer data sets with 70% random dropout. The gray tiles in heatmap indicate missing entries. From left to right: matrix with 70% random dropout, after mean impute, after SVDImpute, after DeepMF. **A** Medulloblastoma data set **B** Leukemia data set **C** TCGA BRCA data set **D** SRBCT data set **E** Clustering accuracy of cancer subtyping on sample latent matrices generated by two imputations and five matrix factorization tools on different cancer data sets with 70% random dropout.

Remainder that alongside the imputation process, DeepMF produced sample latent matrix. To investigate whether missing entries will hinder DeepMF in matrix decomposition, we applied hierarchical clustering into sample latent matrices generated by sparse matrices (**Figure S4**) and computed the clustering accuracy with ground-truth subtyping labels (**Figure 4**). Since the four matrix factorization tools do not accept input with missing values, we fitted the high dimensional matrices treated by MeanImpute and SVDImpute into four baseline approaches, then obtained the corresponding low-dimensional sample latent matrices with rank *K* = 2 for Medulloblastoma data set, rank *K* = 3 for Leukemia data set, rank *K* = 3 for TCGA BRCA data set, and rank *K* = 4 for SRBCT data set, respectively. **Figure 4 E** shows that in terms of clustering accuracy, DeepMF outperforms all eight imputation and factorization combinations, exhibiting the best embedding power with clustering accuracy of 88% for Medulloblastoma data set, 100 % accuracy for TCGA BRCA data set, 84% for Leukemia, and 96% for SRBCT data sets.

## Discussion

In this paper we presented DeepMF, a supervised learning approach to the dimension reduction problem. Unlike current approaches, the method is designed to have high tolerance with respect to noisy data and missing values. Experiments using synthetic and real data corroborated this fact, showing DeepMF to be particularly suited for subtype discovery on omics data.

We have not addressed several issues. The first is with regard to the choice of the three hyper-parameters *K, L, W* in DeepMF. The choice of the reduced dimensionality *K* is arguably difficult, since it is an open problem for the entire dimension reduction research community. Different combination of *K, L* might lead to distinct molecular feature and sample latent matrices. To find the optimal network structure for accurate biological signature interpretation, we defined 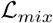 to guide the hyperparameter search. Otherwise, we resort to multiple trials for the tuning of these parameters.

In this paper we have used DeepMF only on mRNA, miRNA, and protein data. However, DeepMF is not limited to these data modality. Human metabolome profiles can certainly benefit from analysis using DeepMF, since the data is known to often suffer from missing values. We intend to apply DeepMF to metabolome and discover signatures beneficial to human health.

In this study, we only utilized the sample latent matrix for subtype detection, we plan to employ molecular feature latent matrix to uncover gene functional pathways in future work.

## Conclusion

MF-based analyses are commonly used in the interpretation of high-throughput biological data. Our proposed DeepMF is an MF-based deep learning framework which overcomes traditional shortcomings such as noise and missing data. Our experiments on simulation data and four omics cancer data sets established DeepMF’s feasibility in denoising, imputation, and in discovering the underlying structure of data.

## Abbreviations

MF: Matrix Factorization
NGS: Next-Generation Sequencing
PCA: Principal Component Analysis
ICA: Independent Component Analysis
NMF: Non-negative Matrix Factorization
ALL: Acute Lymphoblastic Leukemia
AML: Acute Myelogenous Leukemia
TCGA: The Cancer Genome Atlas Network
BRCA: Breast Cancer
SRBCT: Small Round Blue Cell Tumors
BL: Burkitt Lymphoma
EWS: Ewing Sarcoma
NB: Neuroblastoma
RMS: Rhabdomyosarcoma

## Competing interests

The authors declare that they have no competing interests.

## Ethics approval and consent to participate

Not applicable.

## Consent for publication

Not applicable.

## Availability of data and materials

The source code can be found in https://gitlab.deepomics.org/jiaox96/DeepMF

## Funding

Publication costs are funded by the GRF Research Projects 9042348 (CityU 11257316). The work described in this paper was also supported by the project.

## Acknowledgements

We would like to express sincere gratitude to Prof. Yen Kaow Ng (Universiti Tunku Abdul Rahman) for manuscript revision.

## Author’s contributions

S. C. L. conceived the idea.

S. C. L., L. C. designed the network.

L. C, J. X. implemented the network.

L. C, J. X. conducted the analysis.

L. C. drafted the manuscript.

S. C. L. supervised the project, revised the manuscript.

All authors read and approved the final manuscript.

## Supplementary Tables

**Table S1.**
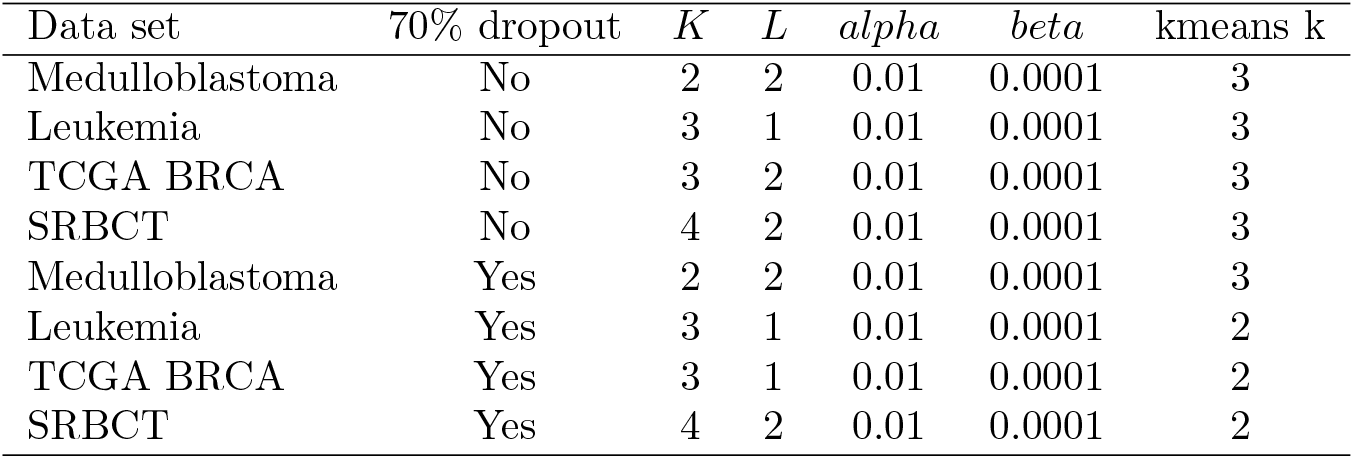
DeepMF training configuration on cancer data sets

| Data set | 70% dropout | $K$ | $L$ | $\alpha$ | $\beta$ | kmeans k |
| --- | --- | --- | --- | --- | --- | --- |
| Medulloblastoma | No | 2 | 2 | 0.01 | 0.0001 | 3 |
| Leukemia | No | 3 | 1 | 0.01 | 0.0001 | 3 |
| TCGA BRCA | No | 3 | 2 | 0.01 | 0.0001 | 3 |
| SRBCT | No | 4 | 2 | 0.01 | 0.0001 | 3 |
| Medulloblastoma | Yes | 2 | 2 | 0.01 | 0.0001 | 3 |
| Leukemia | Yes | 3 | 1 | 0.01 | 0.0001 | 2 |
| TCGA BRCA | Yes | 3 | 1 | 0.01 | 0.0001 | 2 |
| SRBCT | Yes | 4 | 2 | 0.01 | 0.0001 | 2 |

## Supplementary Figures

**Figure S1.**
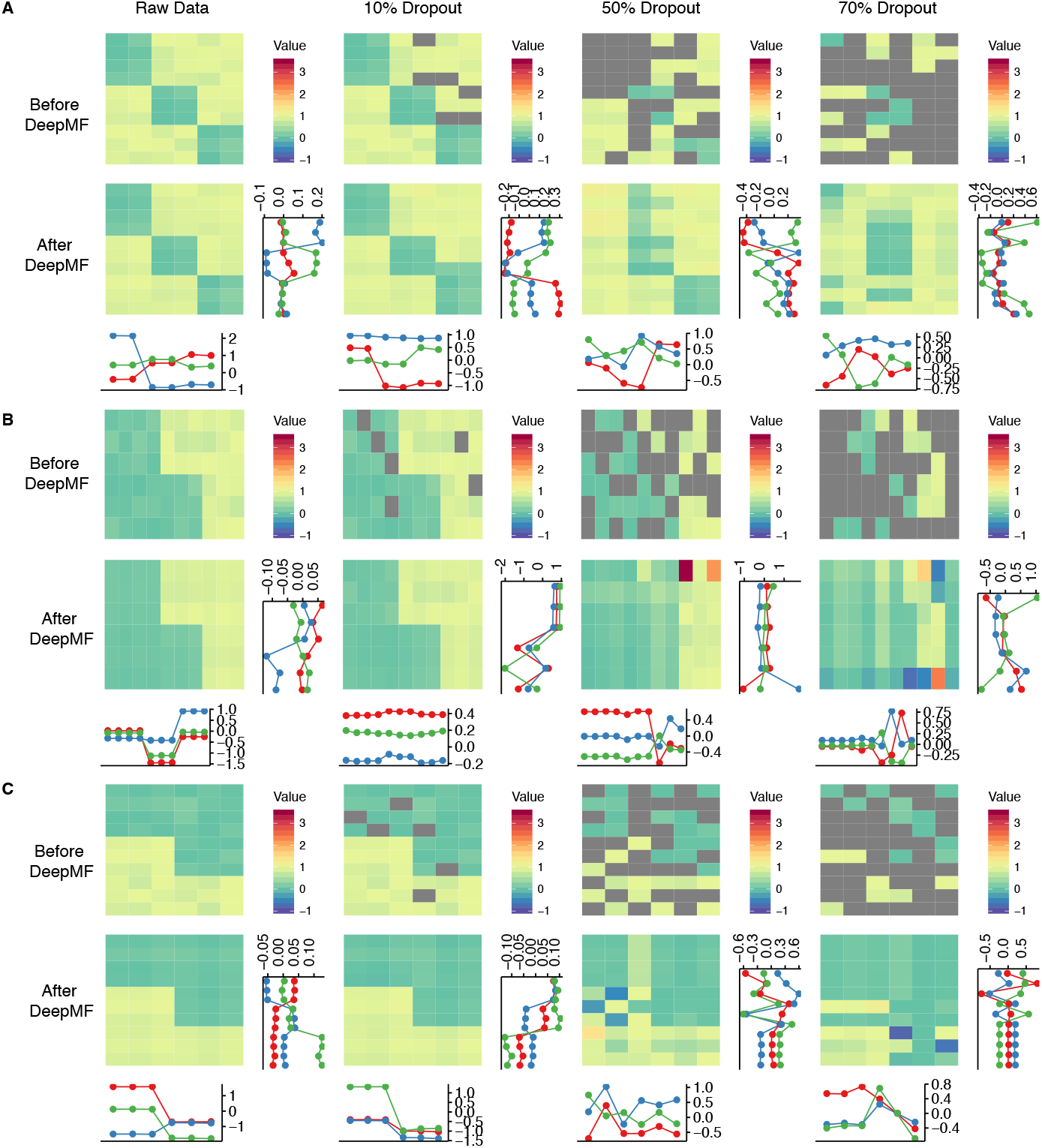
DeepMF performance on 10 × 6 synthetic matrices DeepMF denoising, imputation, and factorization performance on 10 × 6 synthetic matrices with different pattern. Inside each pattern, from left to right: raw matrix, 10% random dropout, 50% random dropout, 70% random dropout; from top to bottom: before DeepMF, and DeepMF. The horizontal line plot show the sample latent factors, the vertical line plot refer to feature latent factors. **A** Matrix with pattern A **B** Matrix with pattern B **C** The transpose matrix of pattern B

**Figure S2.**
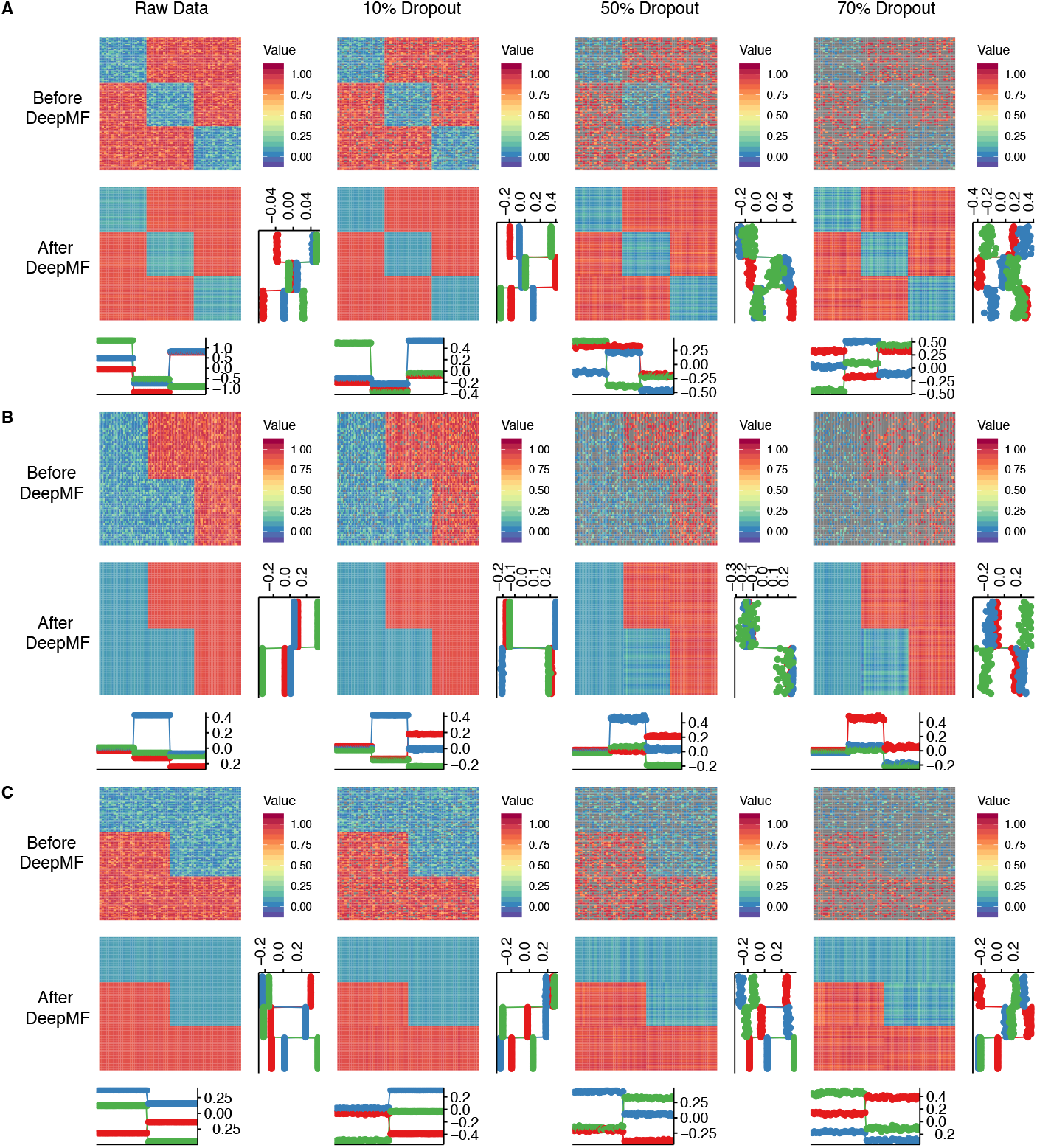
DeepMF performance on 1000 × 600 synthetic matrices DeepMF denoising, imputation, and factorization performance on 1000 × 600 synthetic matrices with different pattern. Inside each pattern, from left to right: raw matrix, 10% random dropout, 50% random dropout, 70% random dropout; from top to bottom: before DeepMF, and DeepMF. The horizontal line plot show the sample latent factors, the vertical line plot refer to feature latent factors. **A** Matrix with pattern A **B** Matrix with pattern B **C** The transpose matrix of pattern B

**Figure S3.**
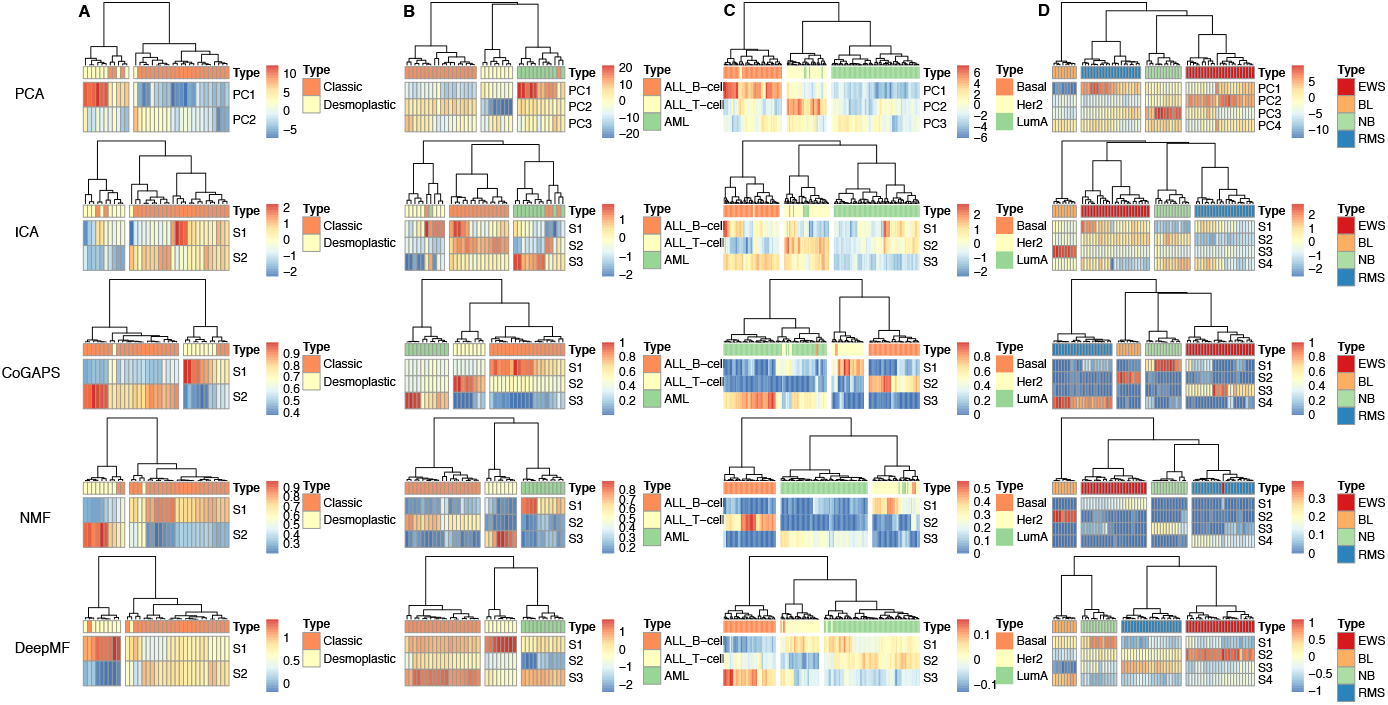
Hierarchical Clustering plot for sample latent matrices Sample latent matrices are generated by five matrix factorization tools on different cancer data sets. From top to bottom, each row represents sample latent matrices generated by PCA, ICA, CoGAPS, NMF, DeepMF. **A** Medulloblastoma data set **B** Leukemia data set **C** TCGA BRCA data set **D** SRBCT data set

**Figure S4.**
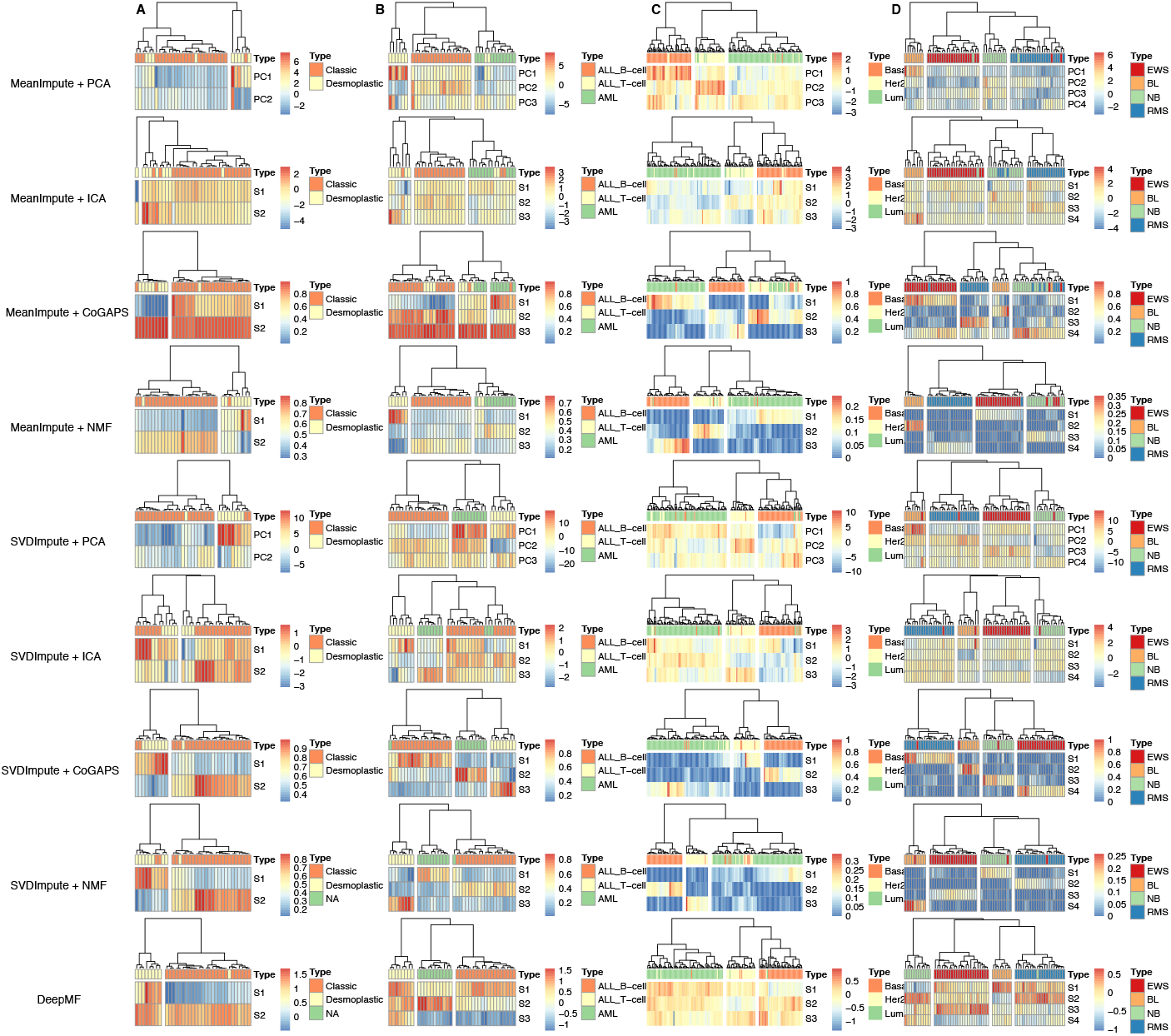
Hierarchical Clustering plot for sample latent matrices generated from 70% random dropout data sets Sample latent matrices are generated by two imputation tools and five matrix factorization tools on different cancer data sets with 70% random dropout. From top to bottom, each row represents sample latent matrices generated by meanImpute + PCA, meanImpute + ICA, meanImpute + CoGAPS, meanImpute + NMF, SVDImpute + PCA, SVDImpute + ICA, SVDImpute + CoGAPS, SVDImpute + NMF, DeepMF. **A** Medulloblastoma data set **B** Leukemia data set **C** TCGA BRCA data set **D** SRBCT data set

